# A core transcriptional response for biofilm formation by *Y. pseudotuberculosis*

**DOI:** 10.1101/2022.03.11.483923

**Authors:** A.K.M.F. Mahmud, K. Nilsson, D.K. Soni, R. Choudhury, R. Navais, Simon Tuck, K. Avican, M. Fällman

## Abstract

Previous transcriptional profiling of the enteropathogen *Yersinia pseudotuberculosis* during persistent stages of colonisation of mouse cecal lymphoid follicles indicated the possible involvement of biofilm in infection maintenance. Not much is known about the mechanisms responsible for biofilm formation by this pathogen, and most current knowledge is based on results of experiments conducted using the related *Y. pestis* pathogen that forms biofilm in the flea gut. In this study, we performed transcriptional profiling of *Y. pseudotuberculosis* in biofilms from different biofilm-inducing conditions, bile exposure, amino acid deprivation and *in vivo* mimicking conditions with and without oxygen. The comparison of differential expression of genes in biofilm versus planktonic bacteria showed a set of 54 core genes that were similarly regulated, independent of inducing condition. This set included many genes that were previously shown to be associated with biofilms, such as *hutG, hsmF, hmsT* and *cpxP* that were upregulated and other genes such as *hmsP* and *rfaH* that were downregulated. There were also novel biofilm-associated genes, including genes encoding hypothetical proteins. To identify the genes involved in inducing biofilm formation, the gene expression of bacteria during an early initial phase when biofilm starts to form after induction by bile or amino acid depletion was determined. Comparisons of the resulting gene expression profiles with the profiles of non-induced bacteria incubated for the same period of time showed a set of core genes associated with early biofilm formation. This set included genes involved in quorum sensing, pili biogenesis and genes indicative of a potential metabolic shift involving nitrogen utilisation. Genes encoding components of sugar phosphotransferase systems were also up-regulated during biofilm induction. Assays of biofilm formation by bacteria deleted of some of these core genes showed that strains lacking *hpr* and *luxS*, which are known to be important for functional sugar phosphotransferase systems and quorum sensing, as well as *glnL* encoding a sensory histidine kinase were most negatively affected. Most of the deletion mutant strains tested were affected, but the effect was less severe, suggesting high levels of redundancy in the pathways involved in biofilm formation by this pathogen.

## Introduction

Bacterial biofilm formation, which is evolutionarily conserved and widespread among pathogenic bacteria, is leading to increasing concerns due to its strong connection to chronic bacterial infections ^1^. Biofilm infections are a major cause of antibiotic overuse in humans as well as in animal husbandry, and contributes to development and spread of antibiotic resistance genes among bacteria ^2^. A major feature of bacterial biofilms is the presence of a self-producing extracellular matrix in biofilms composed of extrapolymeric components, such as polysaccharides, proteins, extracellular DNA, lipids and other biopolymers ^3–5^. Bacteria in biofilms are protected by this matrix and are usually much more resistant to antibiotics; in the infection situation, this results in prolonged antibiotic treatments that are commonly associated with increased morbidity and mortality ^6–8^. Examples of infections involving biofilms are chronic wounds, cystic fibrosis, otitis media, osteomyelitis, prostatitis, chronic pulmonary infections and various nosocomial infections as well as chronic infections caused by bacteria in biofilms on transplanted medical devices ^9–12^.

In recent years, comprehensive studies have been conducted to understand the regulation of biofilm formation in different types of bacteria, for example in *Pseudomonas aeruginosa*, *Salmonella* enterica serovar Typhimurium, *Escherichia coli*, *Vibrio cholera* and *Serratia marcescens* ^13–17^. However, despite many approaches, it is still not clear how biofilms are regulated, and a specific set of core genes that are characteristic of the biofilm state is still lacking ^18^. Studies of different bacteria have shown that pathogenic species can employ different strategies for biofilm formation regarding utilisation of surface structures such as pili, flagella, LPS and exopolymeric substances under specific conditions ^19,20^. The molecular mechanisms that regulate biofilm formation vary among bacterial species, and they involve alternative or multiple pathways, which can also vary between different strains of the same species. Hence, bacteria show great complexity in terms of regulatory pathways for biofilm formation where environmental influences are expected to have an impact. This complicates the development of new drugs that can hinder or weaken bacterial biofilms and contribute to treatment of bacterial infections. Increased understanding of molecular pathways that are important for biofilm formation in different environments is therefore urgently needed.

Enteric *Yersinia* species infect the ileocecal area in humans ^21^, sometimes causing long-lasting persistent infections, which in some cases are associated with development of reactive arthritis ^21–23^. Accordingly, the enteric *Yersinia pseudotuberculosis* YPIII strain was previously shown to be a preferred model organism for studies on persistent bacterial infections in mice where it persists in lymphoid follicles in cecum ^24^. Interestingly, in this context, the persistent stage of the *Y. pseudotuberculosis* infection was found to involve transcriptional reprogramming with upregulation of genes associated with different stress responses, including biofilm-associated genes ^25^. Current knowledge of molecular mechanisms responsible for biofilm formation in *Y. pseudotuberculosis* is however limited. Most of the knowledge about the genes involved comes from studies of its genetically very similar relative, the plague-causing *Yersinia pestis.* Many *Y. pestis* genes known for their biofilm-causing properties are conserved and have been found to have similar functions in *Y. pseudotuberculosis* where, for example, the Hms system has been shown to play a role ^26^. There are, however, also differences between these species. For example, *Y. pestis* can form biofilm in the flea gut, which requires the murine toxin Ymt, that is encoded on its pMT1 plasmid and is not present in *Y. pseudotuberculosis* ^27^. It also forms biofilm in the nematode mouth ^28^, and the *Y. pseudotuberculosis* YPIII strain is one of very few *Y. pseudotuberculosis* strains that can do the same – a property suggested to be a result of the inactivation of the regulatory *phoP* gene ^26^. Further, the genes encoding the biofilm repressors Rcs and NghA are inactive in *Y. pestis* but functional in *Y. pseudotuberculosis* ^27,29^. All these variations suggest the existence of complex regulation of biofilm formation in *Yersinia species*. In addition to the biofilm formation associated with infection of mammalian hosts, the spread of *Y. pseudotuberculosis* through carrots, lettuce and raw milk as well as its environmental maintenance has been suggested to require biofilm formation ^30–33^. Although some biofilm regulatory elements in *Y. pseudotuberculosis* are known, more are to be identified. Additionally, many downstream target genes are yet to be characterised, and therefore, the functional core for the biofilm circuit is yet to be elucidated.

Genome-wide transcriptomic analysis by RNA sequencing (RNA-Seq) is becoming a favoured tool to study regulation in biofilms and the planktonic state of bacteria. Several studies highlight a significant number of genes that are differentially expressed in the biofilm state ^34–39^. However, comparative transcriptome studies of biofilms formed by pathogenic bacteria under different environmental conditions are rare. To fill this gap and reveal biofilm formation circuits in *Y. pseudotuberculosis*, we aimed to identify genes associated with biofilm formation induced by different stresses. We could identify a set of core genes that were similarly regulated in bacteria in mature biofilms independent of the inducing environment. The mature biofilm core genes included both known biofilm regulators and associated genes as well as novel genes not previously reported as biofilm-associated genes. We also identified pathways activated in the early phases of biofilm induction. Potentially interesting genes were mutated, and their importance for biofilm formation was systematically evaluated.

## Results and Discussion

### Influence of environmental factors on biofilm formation by *Y. pseudotuberculosis*

In order to obtain a general view of biofilm formation by *Y. pseudotuberculosis*, we first investigated its ability to form biofilms in different environments. Since this strain of *Y. pseudotuberculosis* has lost the cellulose synthesis operon *bcsQABZC* ^40^, its biofilm was expected to be partly distinct from what is commonly seen, such as in the case of *S.* Typhimurium and *P. aeruginosa*. Cellulose production contributes to a wrinkled phenotype on agar plates, a classical assay for biofilm formation for many bacteria. When comparing the colony phenotype with cellulose producing bacteria such as the related enteropathogens *E. coli, S.* Typhimurium and *P. aeruginosa,* it was obvious that this phenotype was absent in *Y. pseudo-tuberculosis* (Figure 1A). However, the utilisation of other methods such as crystal violet staining of biofilm material on glass tube walls showed that this *Y. pseudotuberculosis* strain indeed formed biofilm (Figure 1B). Electron microscopy of the biofilm showed accumulations of bacteria with some of them growing in chains (Figure 1C). Biofilm formation by *Y. pseudotuberculosis* was also obvious with ECtracer™ ^41^, a fluorescent tracer molecule that binds cellulose and curli in the extracellular matrix (Figure 1D). Moreover, WGA-staining of *Caenorhabditis elegance* larvae exposed to *Y. pseudotuberculosis* showed bacterial biofilm in the larvae mouth (Figure 1E) as previously shown for both *Y. pestis* and *Y. pseudotuberculosis* ^42^.

**Figure 1:**
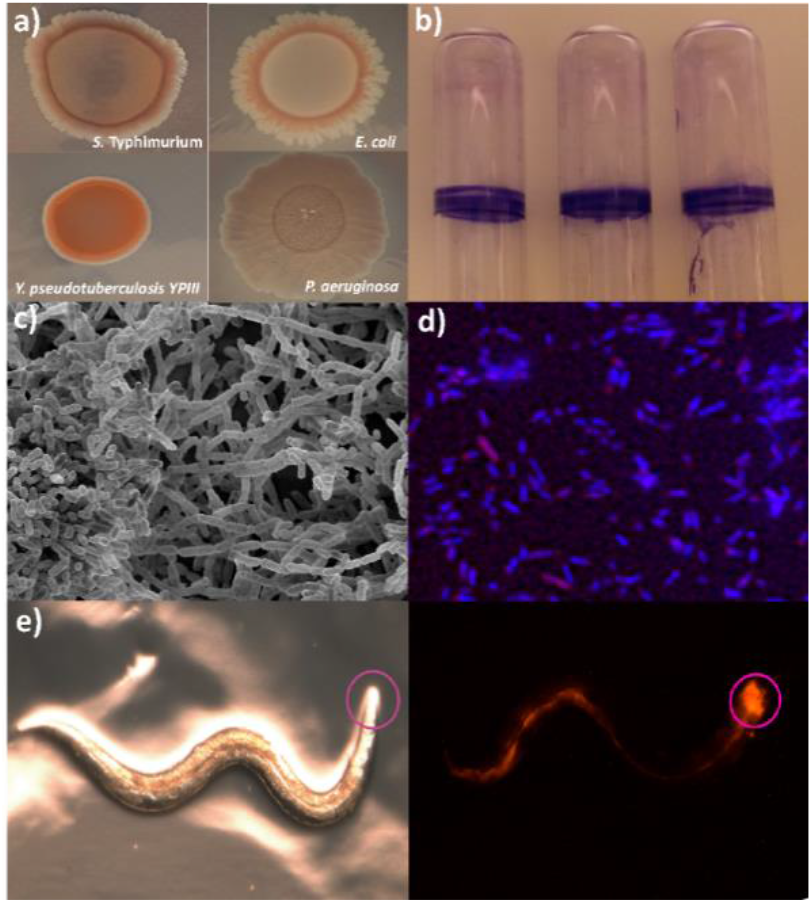
*Y. pseudotuberculosis* biofilm formation. a) *S.* Typhimurium, *E. coli, Y. pseudotuberculosis* YPIII and *P. aueruginosa* cells grown on congo-red agar plates for 4 days. b) *Y. pseudotuberculosis* biofilm formation in glass tubes, stained with crystal violet. C) Scanning electron microscopy picture at 2,500 times magnification of *Y. pseudotuberculosis* biofilm induced by incubation for 48h in LB supplemented with bile d) Fluorescence microscopy of *Y. pseudotuberculosis* biofilm stained with EC-tracer, bacteria ^30^ and extracellular matrix (blue). e) Light stereomicroscope (left) and fluorescence stereomicroscope (right) images of *C. elegans* incubated overnight together with *Y. pseudotuberculosis-*tdTomato (*C.elegans* head is encircled).

Hence, *Y. pseudotuberculosis* forms biofilms on different surfaces and under different conditions. The exact composition of the biofilms is not known, but extracellular DNA, polysaccharides, proteins, and as indicated by the EC-tracer staining also curli materials are likely to contribute.

To determine the most suitable conditions for global transcriptomic studies of biofilm bacteria, we set out to explore conditions and agents influencing *Y. pseudotuberculosis* YPIII biofilm formation. Biofilm formation could be detected during growth at 26°C, but was more prominent at 37°C, both in rich and minimal media (Supplementary Table S1). Applying known inducing conditions, such as the addition of bile or depletion of amino acid (AA-), resulted in increased biofilm formation, which was especially extensive at 37°C. Moreover, the addition of Triton X-100 and growth under anaerobic conditions at 37°C were associated with a high amount of biofilm formation. Some agents or treatments, such as high concentrations of divalent ions, glucose or depletion of iron, had a suppressive effect on biofilm formation (Supplementary Table S1).

To get a broader picture of biofilm formation, we also wanted to include conditions that were more relevant to host infection. For this purpose, we utilised the media used for culturing mammalian cell lines – in this case, MEM supplemented with glutamine, serum and bile. We found that *Y. pseudotuberculosis* also formed biofilm when incubated in this *in vivo* mimicking medium (IVM) at 37°C under an atmosphere of 5% CO_2_, and that it also did so under anaerobic conditions (IVM-An) although to a lower extent (Figure 1; Supplementary Table S1).

### RNA-extractions from biofilm bacteria require optimised protocols

From the initial experiments, bile and AA-as well as IVM and IVM-An were chosen for the planned transcriptomic analyses of *Y. pseudotuberculosis* biofilms. To get data sets from these conditions that were comparable with each other, it was important to determine optimal conditions and time points for sampling. For this purpose, we considered two important points: one was the viability of the biofilm and planktonic bacteria at the time of sampling, and the other was the stability of the biofilm. Viability assays of biofilm bacteria showed that prolonged incubation time, which resulted in an increased amount of biofilm mass, commonly showed a high extent of dead bacteria, which could be rescued by changing media at certain time points (Figure S1A-C). The different conditions required different sampling times to obtain a stable biofilm with viable bacteria. A concentration of 0.5% bile in Luria Broth (LB) at 37°C was found to be suitable to induce high amounts of biofilm with viable bacteria after 48 hours (Figure 2A). For AA-, we aimed for an amino acid depletion condition that did not significantly affect growth but provided stable biofilm. By testing the different combinations of deleted amino acids, we found that depleting three amino acids from different amino acid classes (tyrosine, histidine and tryptophan) resulted in a high amount of biofilm formation after 30 hours at 37°C with a necessary change of media after 18 hours (Figure 2B and Figure S1A). For IVM and IVM-An, it was necessary to change media after 24 hours and sample after 36 hours (Figure 2C C–D and Figure S1B-C).

**Figure 2:**
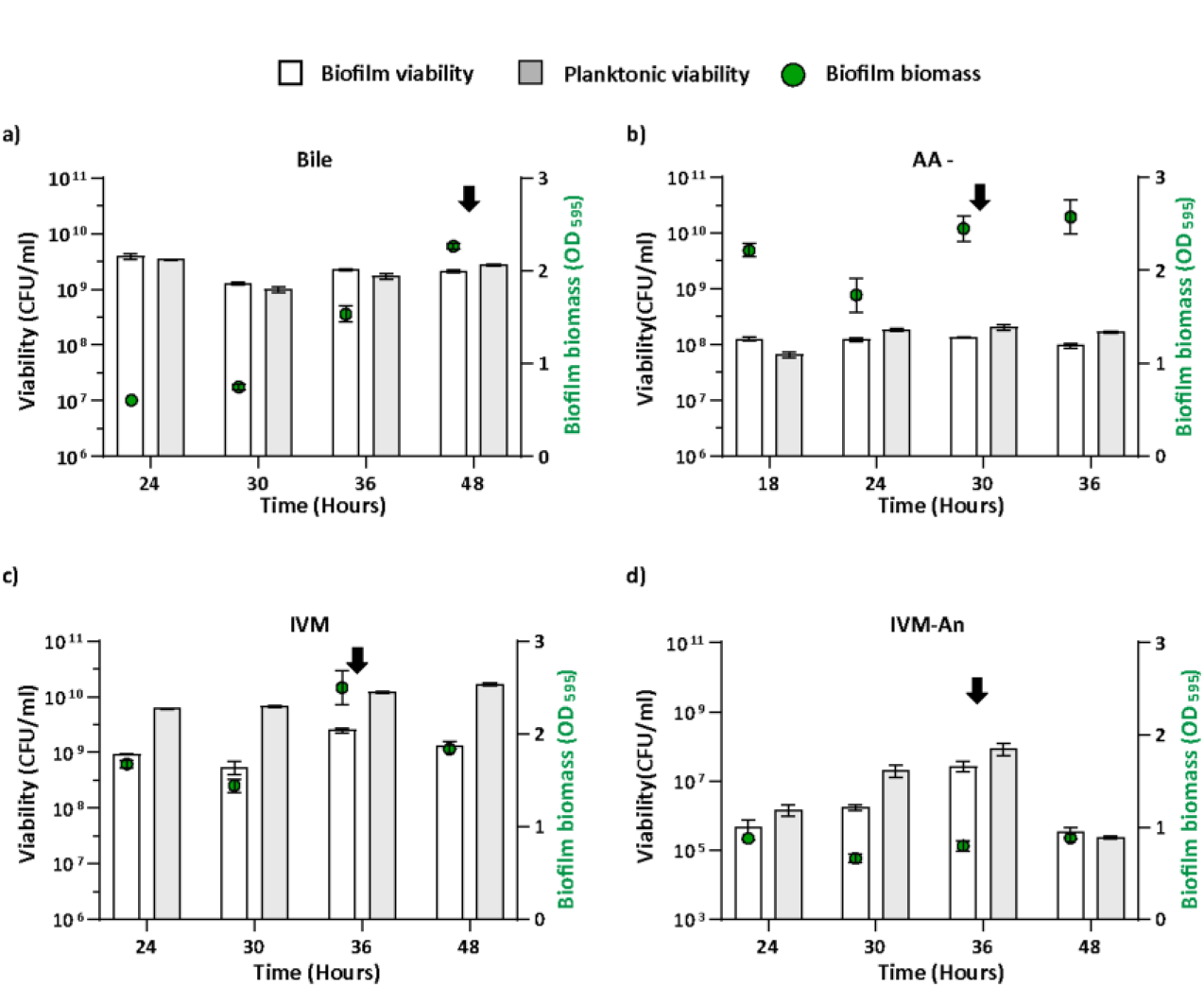
Viability and Biofilm mass measurements as a basis for RNA-extraction. Viable counts of *Y. pseudotuberculosis* biofilm and planktonic bacteria under different conditions at indicated time points (left axis, white and grey bars) and corresponding biofilm biomass (green right axis and points) revealed by Crystal violet staining of biofilm material on glass tube wall. a) biofilm induced by bile; RNA collected at 48 hours. b) biofilm induced by AA-; media changed after 18 hours and RNA collected at 30 hours. c) and d) biofilm formed in IVM and IVM-An; media changed after 24 hours and RNA collected at 36 horus. Three individual biological replicates were used for the viability and biofilm biomass measurement and presented as CFU. per ml +/- SEM and OD595 +/- SEM.

The preparation of bacterial RNA from the collected biofilm material required several modifications and optimisations of the conventional protocol. One important step was to get rid of DNA, indicating extracellular DNA as one major component of the biofilm matrix. For this purpose, we included a step for prolonged DNase treatment in the RNA extraction protocol (see details in “Materials and methods”). It was also obvious that AA- and IVM-An biofilm bacteria required alternative lysis protocols to obtain RNA of good quality.

### Quorum sensing and altered expression of membrane transporters characterise bacteria in mature biofilm

**T**he resulting RNA-Seq data were processed and analysed by ProkSeq, an optimised data analysis pipeline for prokaryotic organisms ^43^, unless otherwise mentioned in materials and methods. Given the heterogeneous nature of biofilms, we put extra emphasis on checking the quality of the data, including filtering to reduce noise due to technical artifacts such as PCR duplicates. This was especially required for the IVM-An data set, which also showed lowest genome coverage (Figure S2). After removing the PCR duplicates, the average library depth of anaerobic bacteria was still in the range of ~ 1.5 million reads per library.

For comparisons of the different data sets, the cut-off for fold change was set to ≥ 1.5 and the P adjusted value to ≤ 0.05. For each condition, a robust transcriptomic change between planktonic and biofilm bacteria was observed (Figure 3A). It was noted that a significant number of genes were not detected in IVM-An, neither from biofilm nor planktonic bacteria. The reason for this can be a suppressed gene expression rather than an effect of too shallow sequencing, which is an assumption supported by the high abundance of PCR duplicates in the raw data set. Nevertheless, differential expression analyses showed that there were many genes that were differentially expressed under this condition, but the fold changes were generally lower (2- to 4-fold) compared to what was seen for bile, AA- and IVM aerobic (Figure 3A).

**Figure 3:**
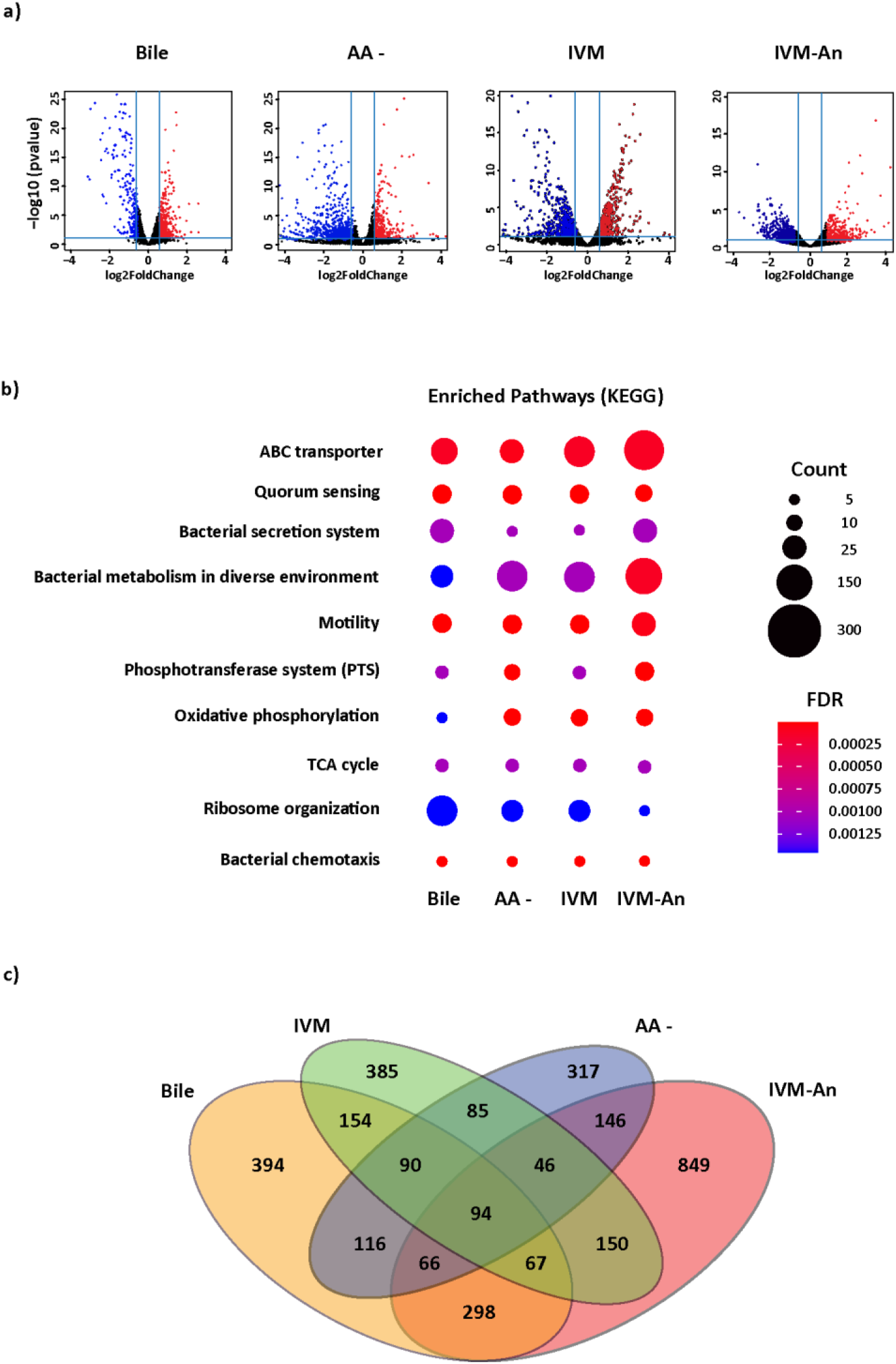
*Y. pseudotuberculosis* differential gene expression in mature biofilm formed under different conditions. a) Volcano plots showing *Y. pseudotuberculosis* differential gene expression patterns (biofilm/planktonic) under the different experimental conditions. Y axis indicates the statistical significance as False discovery rate (FDR) adjusted *P-*value in negative logarithmic scale. X axis indicates the fold change in gene expression (biofilm/planktonic). Each dot represents a gene, red resemble upregulation and blue downregulation of gene expression in biofilm bacteria compared to planktonic. b) Dot plot showing common enriched pathways of differential expressed genes in biofilm bacteria in the tested conditions. Size of the dots indicates the number of genes present in the pathway and colour shows the enrichment score of the pathway in FDR adjusted *P*-value. c) Venn diagram showing the number of differentially expressed genes in the conditions and their overlaps.

A comparison of differential gene expression by bacteria in the different biofilms showed a large variability (Figure 3C and Supplementary Table S2). Given the nature of the gene products of the differentially expressed genes, it was obvious that this, to a great extent, reflected responses to the environment *per se*. This was particularly obvious for samples from bacteria in the anaerobic environment where the number of genes only regulated during this condition was as high as 849. To reveal the eventual common molecular functions that are active in biofilm bacteria under the different conditions, data from differential expression analyses were subjected to KEGG pathway enrichment analyses. These analyses showed 10 KEGG pathways that were enriched in all conditions even though the number of genes found in the pathways and the significance of the enrichment differed between conditions (Figure 3B).

Among the enriched KEGG pathways, ABC transporters, quorum sensing, chemotaxis and bacterial motility were found in all conditions. Genes involved in these pathways have also been observed in other transcriptomic and experimental studies. IVM-An biofilm bacteria showed the highest expression of genes encoding different ABC transporters, which suggested that these bacteria might be less metabolically active compared to bacteria in aerobic biofilms ^44^. This assumption was further supported by the lower expression of genes involved in ribosome organisation, indicative of a lower growth rate of IVM- An biofilm bacteria. Nevertheless, despite high divergence in gene expression between bacteria in the different biofilms, there were common functional signatures, which suggested the existence of key determinants that are critical for the formation of biofilm.

### Core genes for mature biofilm

Although differential expression analyses showed divergence in the gene expression profiles reflecting the different conditions, there was also a set of 94 genes that were differentially regulated between planktonic and biofilm bacteria in all conditions used (Figure 3C). Among these genes, 58 showed similar regulation where the majority (53 genes) were upregulated in biofilm bacteria (Figure 4A). This gene group was named “core genes for mature biofilm” (Table 1). Some of the genes in this group that were upregulated in biofilm bacteria, such as *hmsF, hmsT, cpxA* and *cpxP*, are known regulators of biofilm, and some other genes, such as genes involved in curli/pili production and c-di-GMP signalling, have also been associated with biofilms. However, there were many other genes that have not been previously associated with biofilm, including 14 genes encoding hypothetical proteins whose functions are yet to be characterised (Figure 4A and Table 1). Further, among the four genes that were downregulated in biofilm bacteria, there were genes that have been previously associated with the negative regulation of biofilms, such as *hms*P and *rfaH* ^45,46^.

**Figure 4:**
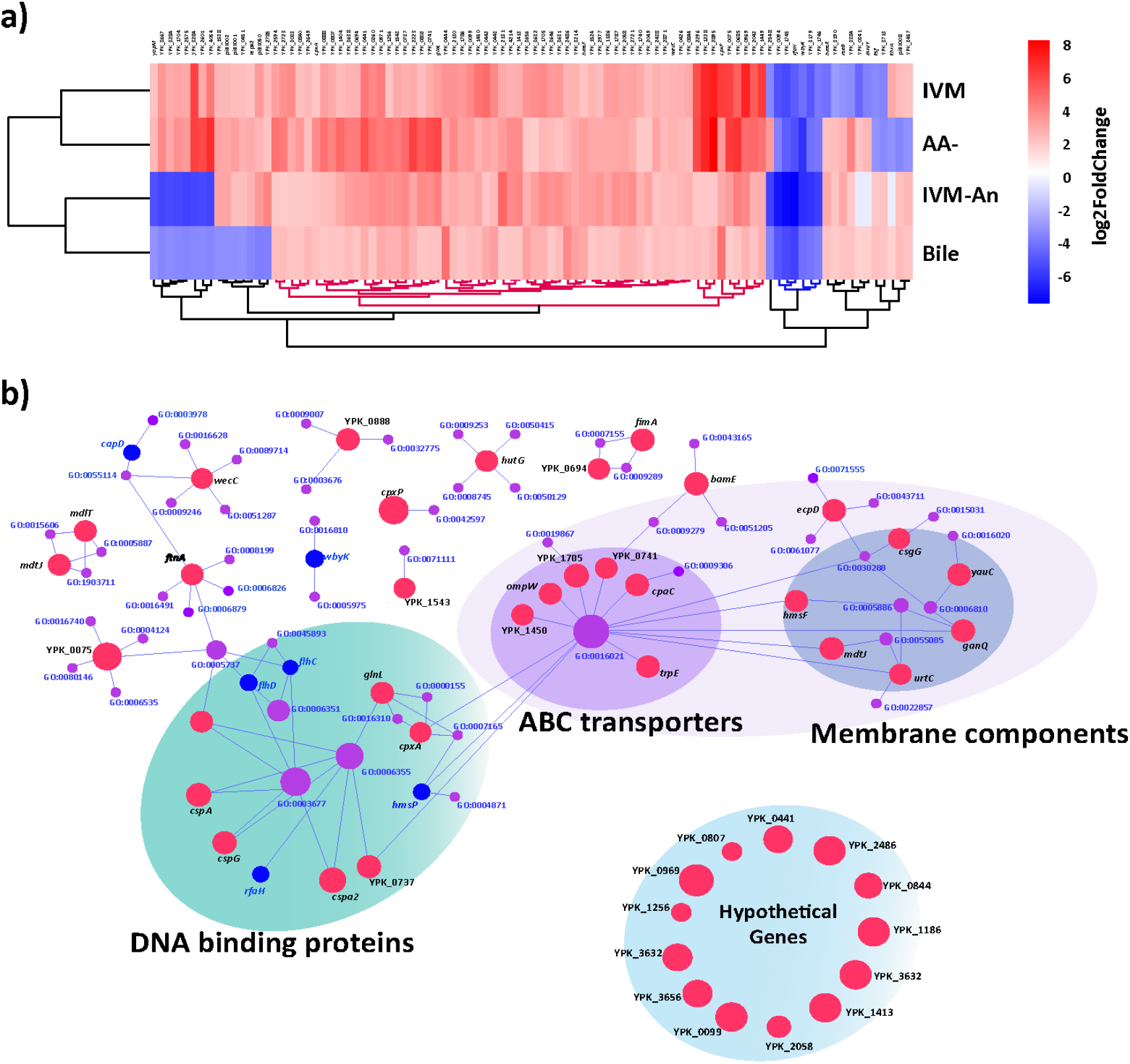
Core genes of mature biofilm. a) Heat maps showing commonly differentially expressed genes (biofilm/planktonic) in the different conditions. Colors indicate fold changes (red upregulation and blue downregulation) of genes in biofilm bacteria. Hierarchical clustering (bottom) was done to cluster genes with similar expression pattern. b) Gene ontology (GO) interaction network created by Cystoscope. Size of the GO term (purple color) is based on number of interacting genes. Red and blue filled circles indicate upregulation versus downregulation of the genes in biofilm bacteria and the size corresponds to differential expression in fold change. Genes encoding hypothetical genes (not represented by GO terms), have been manually added

Due to the limited number of genes, Gene Ontology (GO) network enrichments were used instead of KEGG pathway enrichments to reveal functional linkages between the core genes for mature biofilms. This analysis showed two distinct clusters of genes: one with genes encoding DNA binding proteins including many transcription factors and another with genes encoding membrane-associated proteins with a distinct subgroup of ABC transporters (Figure 4B).

In addition to these, there were multiple genes encoding proteins not connected to one particular GO class; instead, many of the genes belonged to multiple GO classes, suggesting functional diversity (Figure 4B). These core genes were also subjected to analysis using experimentally validated protein-protein interaction data from the string database ^47^. However, this did not show any obvious clusters, but some of the core gene products could interact with each other, or with multiple proteins (Figure S3). Hence, these core genes might represent genes from different pathways, thereby reflecting a diversity of pathways involved in biofilm formation.

### Crucial genes for early biofilm

The 58 core genes identified for bacteria in mature biofilms do not necessarily play important roles in the early phase of biofilm formation where bacteria encounter the initial stress, which leads to the activation of the pathways that are essential for the initiating formation of biofilm. It was not obvious how to identify such genes but given that bile and AA- induce more biofilm compared to non-induced bacteria when incubated at 37°C (Figure 1 and Supplementary Table 1), it should be possible to distinguish between these two groups at the transcriptional level. Further, since these two conditions involve different initial stresses, with envelop stress for bile ^48,49^ and stringent responses for AA- ^50^, these biofilm-inducing conditions can allow determination of core determinants involved in the initiation of biofilm formation independent of the initial stress. Therefore, we first analysed the time point at which the biofilm could be detected for bacteria induced by bile or AA-, while there was none or neglectable amounts of biofilm formed by non-induced control bacteria. Induction by bile and AA- were found to follow a similar pattern during the first hours where biofilm formation started to be detectable after 2– 3 h, while almost no biofilm was seen for non-induced bacteria (Figure 5A).

**Figure 5:**
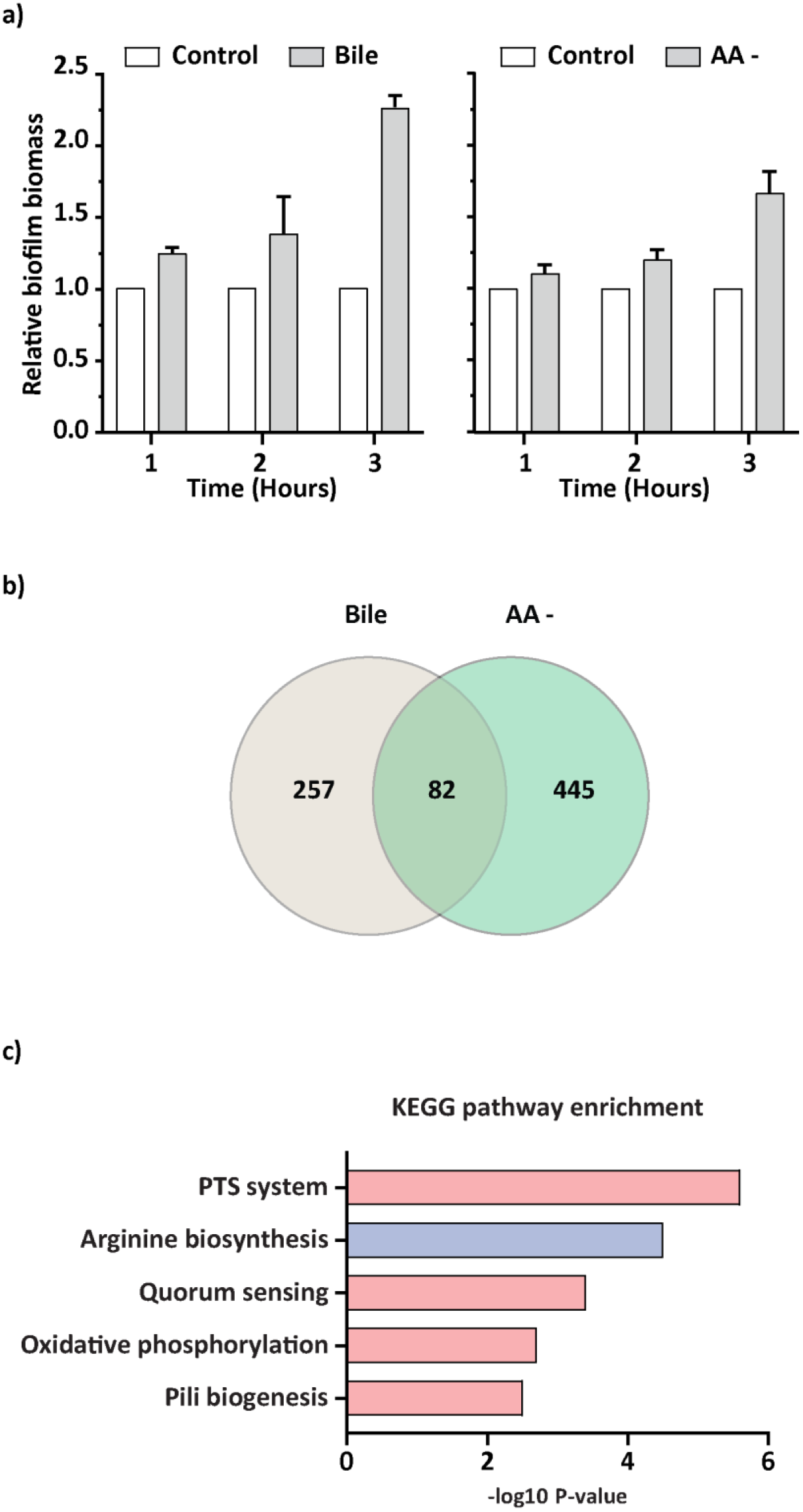
Core genes of early biofilm formation. a) Biofilm biomass measurement of biofilms induced by bile and AA-. Each experiment was conducted in three biological replicates and presented as relative biofilm biomass compared to biofilm formation by untreated bacteria +/- SEM. b) Venn diagram showing number of differentially expressed genes (treatment/control) in bacteria in biofilms induced by bile and AA-, and their overlap. c) Bar graph showing KEGG pathway enrichments among genes that were differentially expressed (upregulated pathways in red and downregulated in blue) by bacteria in both bile and AA- biofilms. Pathways are ranked according to enrichment score in FDR adjusted P-value (X axis).

**Figure 6:**
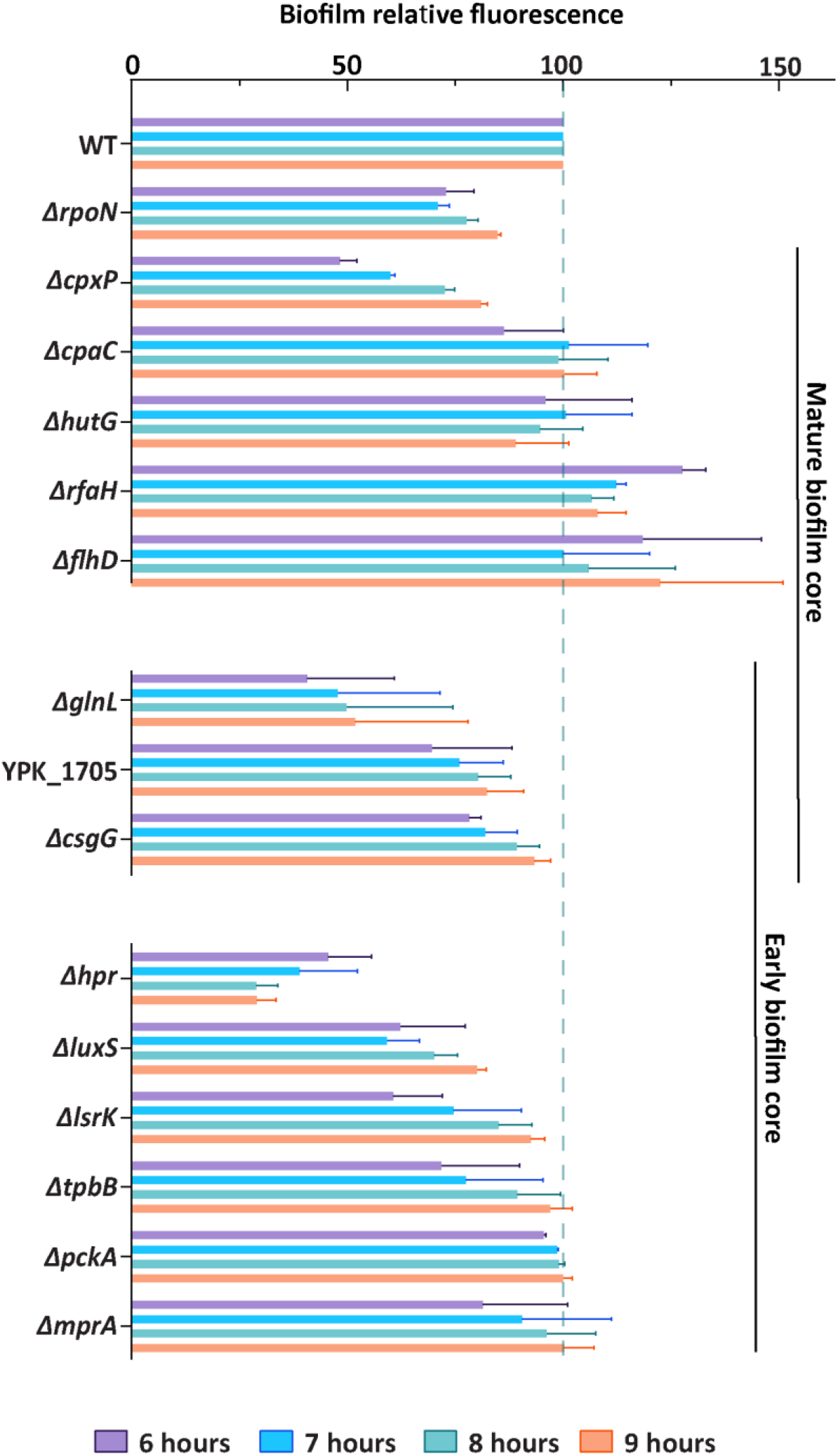
Many core gene products are required for efficient biofilm formation by *Y. pseudotuberculosis*. Biofilm formation by strains deleted of selected biofilm core genes measured by an ECtracer™ biofilm assay after indicated incubation times at 37°C. The different deletion strains lack mature biofilm core genes (upper panel), early biofilm core genes (lower panel), or genes found in both cores (middle panel). All experiments were done in three independent biological replicates with three technical replicates for each. Error bars were calculated according to standard error of the mean (SEM).

These results indicated that it should be possible to extract information about early biofilm-associated gene expression by comparing the global gene expression of bacteria induced using bile or AA-for 3 hours with that of non-induced control bacteria.

Analyses of differential gene expressions by comparing the transcriptional profiles of control and biofilm-induced bacteria (planktonic and biofilm bacteria recovered together) revealed a set of genes distinctly regulated by bile or AA- induction. For bile induction, there were more than 300 differentially expressed genes, and for AA-, there were more than 500 differentially expressed genes (Figure 5B, Supplementary Table S3). Comparisons of the two groups showed that more than 80 genes were commonly regulated in the two conditions (Table 2). In this set of early biofilm core genes, only seven were found among the group of core genes for mature biofilm. The reason might be that core genes for mature biofilm may not be expressed during the early stages of biofilm formation but can instead be required for the maintenance of the biofilm. Further, in accordance with what was shown for the core genes for mature biofilm (but not the same genes), there were genes encoding proteins involved in pili/curli biogenesis and c-di-GMP signalling in this group of core genes commonly regulated at the early phase. The upregulation of pili/curli-associated genes was more pronounced compared to that in the mature phase, suggesting roles of these gene products both in adhesion and extracellular matrix formation. Interestingly, in addition to the genes involved in c-di-GMP signalling, a gene encoding the ABC transporter PstA, previously reported to constitute the receptor for c-di-AMP was upregulated ^51,52^. This bacterial second messenger system have been implicated to be involved in sensing DNA-integrity, cell wall metabolism, fatty acid synthesis and potassium transport ^51^. Other genes of the same operon, *pstB* and *pstC,* that together with *pstA* make up a phosphate transfer system responding to low levels of phosphate, were also found in this core group. Additionally, quorum sensing genes encoding proteins involved in the production of autoinducer-2, were in this core group. Quorum sensing has previously been shown to be important for biofilm formation, as it plays a role in the initiation phase in particular, which requires the presence of a certain amount of bacteria. Further, also in this group, there were many genes encoding hypothetical proteins whose roles are yet to be determined.

### Initiation of biofilm formation involves induction of the sugar phosphotransferase system and quorum sensing and is associated with a potential metabolic switch

Analyses of potential functions of the genes in the group of early biofilm core genes by utilising KEGG pathway enrichments showed five distinct KEGG pathways that were overrepresented (Figure 5C). These included the quorum sensing and pili biogenesis pathways as well as oxidative phosphorylation and the sugar phosphotransferase system (PTS) that were all upregulated, whereas the arginine biosynthesis pathway was downregulated in biofilm bacteria. The upregulation of sugar PTS systems that involve the utilisation of sugar, has previously been shown to play an important role in biofilm formation ^51–54^. Among the enriched pili biogenesis genes, most were connected to the type IV class family that involves curli production, which can contribute to adhesion and microcolony formation ^55–60^. Type IV pili also contribute to flagellar independent motility, the so-called twitching motility, which has previously been shown to be important for bacterial attachment to liquid surfaces in association with biofilm formation ^58,59^. The observed upregulation of oxidative phosphorylation activities may be due to a general adaptive stress response or play a role in metabolic reprogramming for biofilm development as shown in *Bacillus subtilis* ^61^. In accordance with the latter, the observed downregulation of arginine biosynthesis can indicate metabolic reprogramming. By preventing arginine biosynthesis, bacteria can increase catabolism using the nitrogenous bases either for the TCA cycle or for the reciprocal regulation to prevent a futile cycle.

Many of the core genes identified for *Y. pseudotuberculosis* in early and mature biofilm stages have previously been shown to be involved in biofilm formation, whereas other genes are, potentially, novel biofilm genes (Table 1 and 2). To reveal their eventual importance for biofilm formation by this pathogen, some of the genes were deleted and the resulting deletion mutant strains were tested for their ability to form the biofilm. The genes selected represented genes known to be involved in biofilm formation in other bacteria as well as novel genes (Table 1 and 2), including genes involved in c-di-GMP signalling, sugar PTS and quorum sensing. Due to the presence of autoinducer genes in this core group, we also created a Δ*luxS* mutant strain to evaluate the importance of the main autoinducer regulator. Biofilm measurements using ECtracer showed that most of the mutant strains had reduced capacity to form biofilm compared to that of the parental strain (Figure 5 and Tables 1 and 2). However, for both core groups, most of the mutant strains formed some level of biofilm, and with time, many could catch up to the level seen for the parental strain. This suggested a high level of redundancy among pathways or mechanisms contributing to biofilm formation in bacteria, which was not surprising considering the expected fundamental importance of this function in general.

For core genes of the mature biofilm, the strain mutated in a gene encoding a hypothetical protein, YPK_1705, had a demonstrably reduced ability to form a biofilm. This protein is a putative surface protein with a glycin zipper domain, potentially leading to fibril formation, and may contribute to adhesion and the extracellular matrix. Reduced biofilm formation was also seen for strains lacking genes encoding the periplasmic stress response protein CpxP and the curli assembly/transport component CsgG, both of which have been previously shown to be important for biofilm formation in other bacteria. For the early biofilm core genes, two mutant strains were more severely affected compared to the others, Δ*hpr* and Δ*luxS*, both representing genes critical for two functions identified in the functional analyses, namely sugar PTS and quorum sensing. Accordingly, another strain mutated in a gene involved in quorum sensing, *lsrK,* encoding the autoinducer kinase did also show reduced ability to form biofilm. Reduced biofilm formation was also seen for the strains lacking *tpbB*, a gene encoding a di-guanylate cyclase, as well as *mprA* encoding a transcriptional repressor.

Taken together, our data show that biofilm formation in *Y. pseudotuberculosis* is complex and involves many pathways that can contribute where c-di-GMP and, possibly, also c-di-AMP signalling are involved and that PTS and quorum sensing systems have a central role in initiation of biofilm formation. It is also clear that bacteria in biofilm adapt their metabolism and modulate it according to the niche.

## Materials and methods

### Bacterial strains and growth conditions

Strains and plasmids are listed in Table 3. The *Yersinia pseudotuberculosis* YPIII strain was used for evaluation of its biofilm formation in this study. *Escherichia coli* 17-1 λpir was used for cloning and conjugation. Antibiotics were used in the following concentrations: ampicillin (100 μg/ml), kanamycin (50 μg/ml) and chloramphenicol (25 μg/ml). All strains were routinely grown at 26°C in LB medium containing kanamycin (Km; 50 μg/ml). The media for biofilm formation were LB +/- 0.5% bile, Rich MOPS or MOPS depleted of the amino acids tyrosine, histidine and tryptophan (AA-). The IVM medium consisted of MEM-Hepes with glucose supplemented with 10% heat inactivated FCS, 200mM glutamine, and 5% sodium choleate (Sigma).

### Strain construction

In-frame gene deletion mutations in *Y. pseudotuberculosis* were constructed using the In-fusion HD cloning kit (Clontech) according to the manufacturer’s instructions. Briefly, the flanking regions of the respective genes were amplified by PCR and cloned into the suicide vector pDM4 ^62^. This construct was used to transform S17-1 and then transferred into recipient strains through conjugation. Conjugants were purified and incubated on 5% sucrose to recombine out the vector together with the wild-type sequence. The resulting deletions were confirmed by sequencing. Primers used in this study are listed in Table S4.

### Biofilm morphology (Congo red Assay)

Congo red indicator agar plates were prepared by adding 20 mg/ml Congo red (Sigma) and 10 mg/ml Coomassie brilliant blue G (Sigma) to LB agar plates. An aliquot of 100 μl of overnight cultures were plated, and colony morphology was observed after three days of incubation at 26°C.

### Crystal Violet *in vitro* biofilm assay and viability measurements

Biofilm formation was determined as previously described ^63^. The overnight culture was diluted to an OD_600_ of 0.05 and grown to an OD_600_ of 0.5 at 26°C. A 1 ml aliquot of the bacterial culture was pelleted and resuspended in 2 ml of fresh LB media. The suspension was transferred to glass tubes and incubated for at 37°C without shaking for different periods of time. After the incubation, the bacterial suspension was discarded and the tubes were gently washed three times with PBS and stained with 0.1% crystal violet (Sigma-Aldrich) for 15 minutes, followed by successive washing with PBS. Thereafter, the biofilms on the tube surface were dissolved in 33% acetic acid for 15 minutes, and the absorbance at 590 nm was measured with an Ultrospec 2100 Pro Spectrophotometer (Amersham Biosciences).

For viability assays overnight cultures in LB were diluted and grown at 26°C from an OD_600_ of 0.05 to an OD_600_ of 0.5 after which bacteria were pelleted and resuspended in the indicated media. Aliquotes of 5 ml in triplicates were dispensed into tilted Petri dishes and incubated at the indicated conditions. The medium was collected at indicated time points, then transferred to serial dilutions and plated on LB+Km agar plates. The remaining biofilm was washed three times with PBS, detached with a cell scraper, resuspended in 1 ml of PBS and after serial dilutions plated on LB+Km agar plates. Viable cells were counted after 2 days of incubation.

### Scanning electron microscope assay

Samples of *Y. pseudotuberculosis* biofilm were prepared for scanning electron microscopy. Overnight cultures were diluted to an OD_600_ of 0.05 and grown at 26°C to an OD_600_ of 0.8. The cultures were harvested by centrifuging for 5 minutes at 12,000 rpm and resuspended in LB and transferred to wells of 12-well plate containing glass coverslips coated with polylysine (Sigma-Aldrich). The plate was incubated at 37°C for 48 hours after which the biofilm specimens were prepared for electron microscopy as described by Ochmcke et al. ^64^ with some modifications. Biofilm clots were fixed overnight with 2.5% glutaraldehyde. Samples were washed three times with 0.1 M sodium phosphate buffer (pH 7.3), dehydrated with a series of increasing ethanol concentrations (5 minutes in 30%, 5 minutes in 50%, 10 minutes in 70%, 10 minutes in 90% and two times for 10 minutes in 96% ethanol). The dehydrated samples were dried with CO_2_ using the critical point method with an Emitech dryer as outlined by the manufacturer. The dried samples were covered with gold to a 10 nm layer and scanned with a Zeiss DSM 960A electron microscope.

### Staining of biofilm on coverslips with ECtracer

For staining of biofilm with ECtracer™680 (EBBA biotec), 2 coverslips (12×12 mm) were attached with agarose to the bottom and inclining the walls of one well of a 24-well plate. Overnight cultures in LB+Km were diluted in to an OD_600_ of 0.1 and grown at 26°C until an OD_600_ of 0.8. Thereafter, bacteria were pelleted and re-suspended in MOPS and 0.6 ml aliquots of the suspension were transferred to the wells and incubated further at 37°C for various periods of time. The coverslips were thereafter rinsed in PBS and wiped on the backside with a cotton swab. The biofilm on the coverslips was fixed in 4% PFA for 30 minutes, washed in 70% ethanol and then in 30% ethanol for 5 minutes each, then stained for 30 minutes with ECtracer (1000x dilution in PBS) and counterstained with DAPI. The coverslips were mounted on glass slides, and the biofilms were analysed using a fluorescent microscope at 100x magnification using DAPI and TRITC filters.

### Spectrophotometric monitoring of biofilm formation in bacterial cultures with ECtracer

Overnight bacterial cultures grown at 26°C in LB +Km were diluted to an OD_600_ of 0.1 and grown for an additional period of 4h. Samples equivalent to an OD_600_ of 0.8 were pelleted and resuspended in 1ml MOPS + Km. Aliquots of 320ul were mixed with 4.8 μl of 10x diluted ECtracer™680 (EBBA biotec) and thereafter 100 μL were dispensed to a Greiner Flat Black 96-well plate in triplicat-es. The assay was run in kinetic mode at 37°C in a Tecan SparkControle reader using a humidity cassette. Data were collected every hour in Fluorescence Top Reading mode with Excitation wavelength 535 nm, Excitation bandwidth 25 nm, Emission wavelength 680nm, Emission bandwidth 30 nm, and a Gain of 65.

### *In vivo* biofilm assay on *C. elegans*

Nematode growth media agar plates were seeded with the *Y. pseudotuberculosis* YPIII strain expressing the tdTomato gene. The plates were incubated overnight at room temperature (RT). L4 larvae and young adults of *C. elegans* (15–20 individuals) were transferred from Nematode growth media agar plates seeded with *E. coli* OP50 strain to the *Yersinia* seeded plates and incubated overnight at room temperature. Thereafter, the worms were picked from the plates and washed three times with Milli-Q water to remove unattached bacteria. A fluorescence stereoscope (Nikon SMZ800 stereomicroscope) was used to observe the worms and identify attached fluorescent bacteria.

### RNA sequencing and data analysis

For isolation and purification of total RNA from *Y. pseudotuberculosis* biofilm and planktonic bacteria the Trizol method (Ambion Life Technol-ogies, Carlsbad, CA) and the Direct-zol RNA kit (Zymo) were used. For the AA- samples, cell lysis was done using a bead beater and for the IVM-An samples, lysozyme was used. To remove DNA, samples of biofilm bacteria were treated with DNAse in column for 30 minutes at room temperature, except samples from AA- induced biofilms, which were subjected to 2×30 min column DNAse treatments at 37°C. RNA quantity was measured by Cubit (Nordic biolab).

For sequencing, cDNA libraries were prepared using the ScriptSeq™ Complete Kit (Epicentre, Madison, WI, USA) according to the manu-facturer’s instructions. Ribosomal RNA was depleted from total RNA using Ribo-Zero rRNA Removal Kit for Bacteria (Epicenter, Madison, WI, USA) following the manufacturer’s protocol. The resulting cDNA libraries were purified using AMPure XP and quantified using an Agilent 2100 Bioanalyzer. Sequencing was done by Illumina MiSeq and Nova-Seq. For bioinformatic analysis, Prokseq ^43^, a complete RNA-Seq data analysis package was used for data processing, quality control and visualization and differential expression analysis. ProkSeq includes all the tools mentioned below required for data analysis. Reads were aligned with a reference genome *of Y. pseudotuberculosis* YPIII chromosome (NC_010465) and plasmid (NC_006153) using salmon ^65^. Differential gene expression was determined using RUV-Seq ^66^ using an *in silico* predicted control set of genes by in house computer program.

## Data availability

The RNA-seq data files have been deposited in Gene Expression Omnibus under accession number GSExxx. All the computer code and pipeline used in these studies are available on request

## ACKNOWLEDGEMENTS

The work has been supported by funding from Knut and Alice Wallenberg foundation (2016.0063), Swedish research Council (2018-02855), and the Medical faculty at Umea University.

## References

1 Conlon, B. P., Rowe, S. E. & Lewis, K. Persister cells in biofilm associated infections. Adv Exp Med Biol 831, 1–9, doi:10.1007/978-3-319-09782-4_1 (2015).

2 O’Toole, G., Kaplan, H. B. & Kolter, R. Biofilm formation as microbial development. Annu Rev Microbiol 54, 49–79, doi:10.1146/annurev.micro.54.1.49 (2000).

3 O’Toole, G. A. To build a biofilm. J Bacteriol 185, 2687–2689, doi:10.1128/jb.185.9.2687-2689.2003 (2003).

4 Webb, J. S., Givskov, M. & Kjelleberg, S. Bacterial biofilms: prokaryotic adventures in multicellularity. Curr Opin Microbiol 6, 578–585, doi:10.1016/j.mib.2003.10.014 (2003).

5 Battin, T. J., Besemer, K., Bengtsson, M. M., Romani, A. M. & Packmann, A. I. The ecology and biogeochemistry of stream biofilms. Nat Rev Microbiol 14, 251–263, doi:10.1038/nrmicro.2016.15 (2016).

6 Foster, T. J., Geoghegan, J. A., Ganesh, V. K. & Hook, M. Adhesion, invasion and evasion: the many functions of the surface proteins of Staphylococcus aureus. Nat Rev Microbiol 12, 49–62, doi:10.1038/nrmicro3161 (2014).

7 Moskowitz, S. M., Foster, J. M., Emerson, J. & Burns, J. L. Clinically feasible biofilm susceptibility assay for isolates of Pseudomonas aeruginosa from patients with cystic fibrosis. J Clin Microbiol 42, 1915–1922, doi:10.1128/jcm.42.5.1915-1922.2004 (2004).

8 Parsek, M. R. & Singh, P. K. Bacterial biofilms: an emerging link to disease pathogenesis. Annu Rev Microbiol 57, 677–701, doi:10.1146/annurev.micro.57.030502.090720 (2003).

9 Mihai, M. M. et al. Microbial biofilms: impact on the pathogenesis of periodontitis, cystic fibrosis, chronic wounds and medical device-related infections. Curr Top Med Chem 15, 1552–1576, doi:10.2174/1568026615666150414123800 (2015).

10 Del Pozo, J. L. Biofilm-related disease. Expert Rev Anti Infect Ther 16, 51–65, doi:10.1080/14787210.2018.1417036 (2018).

11 Srivastava, S. & Bhargava, A. Biofilms and human health. Biotechnol Lett 38, 1–22, doi:10.1007/s10529-015-1960-8 (2016).

12 Vestby, L. K., Gronseth, T., Simm, R. & Nesse, L. L. Bacterial Biofilm and its Role in the Pathogenesis of Disease. Antibiotics (Basel) 9, doi:10.3390/antibiotics9020059 (2020).

13 Drenkard, E. & Ausubel, F. M. Pseudomonas biofilm formation and antibiotic resistance are linked to phenotypic variation. Nature 416, 740–743, doi:10.1038/416740a (2002).

14 Davey, M. E. & O’Toole G, A. Microbial biofilms: from ecology to molecular genetics. Microbiol Mol Biol Rev 64, 847–867, doi:10.1128/mmbr.64.4.847-867.2000 (2000).

15 Bayramoglu, B., Toubiana, D. & Gillor, O. Genome-wide transcription profiling of aerobic and anaerobic Escherichia coli biofilm and planktonic cultures. FEMS Microbiol Lett 364, doi:10.1093/femsle/fnx006 (2017).

16 Cui, L. et al. CRISPR-cas3 of Salmonella Upregulates Bacterial Biofilm Formation and Virulence to Host Cells by Targeting Quorum-Sensing Systems. Pathogens 9, doi:10.3390/pathogens9010053 (2020).

17 Wood, T. L. et al. Rhamnolipids from Pseudomonas aeruginosa disperse the biofilms of sulfate-reducing bacteria. NPJ Biofilms Microbiomes 4, 22, doi:10.1038/s41522-018-0066-1 (2018).

18 Patell, S. et al. Comparative microarray analysis reveals that the core biofilm-associated transcriptome of Pseudomonas aeruginosa comprises relatively few genes. Environ Microbiol Rep 2, 440–448, doi:10.1111/j.1758-2229.2010.00158.x (2010).

19 Mah, T. F. et al. A genetic basis for Pseudomonas aeruginosa biofilm antibiotic resistance. Nature 426, 306–310, doi:10.1038/nature02122 (2003).

20 Bjarnsholt, T. et al. Pseudomonas aeruginosa tolerance to tobramycin, hydrogen peroxide and polymorphonuclear leukocytes is quorum-sensing dependent. Microbiology (Reading) 151, 373–383, doi:10.1099/mic.0.27463-0 (2005).

21 Puylaert, J. B., Van der Zant, F. M. & Mutsaers, J. A. Infectious ileocecitis caused by Yersinia, Campylobacter, and Salmonella: clinical, radiological and US findings. Eur Radiol 7, 3–9, doi:10.1007/s003300050098 (1997).

22 Hoogkamp-Korstanje, J. A., de Koning, J. & Heesemann, J. Persistence of Yersinia enterocolitica in man. Infection 16, 81–85, doi:10.1007/BF01644307 (1988).

23 Galindo, C. L., Rosenzweig, J. A., Kirtley, M. L. & Chopra, A. K. Pathogenesis of Y. enterocolitica and Y. pseudotuberculosis in Human Yersiniosis. J Pathog 2011, 182051, doi:10.4061/2011/182051 (2011).

24 Fahlgren, A., Avican, K., Westermark, L., Nordfelth, R. & Fallman, M. Colonization of cecum is important for development of persistent infection by Yersinia pseudotuberculosis. Infect Immun 82, 3471–3482, doi:10.1128/IAI.01793-14 (2014).

25 Avican, K. et al. Reprogramming of Yersinia from virulent to persistent mode revealed by complex in vivo RNA-seq analysis. PLoS Pathog 11, e1004600, doi:10.1371/journal.ppat.1004600 (2015).

26 Sun, Y. C., Koumoutsi, A. & Darby, C. The response regulator PhoP negatively regulates Yersinia pseudotuberculosis and Yersinia pestis biofilms. FEMS Microbiol Lett 290, 85–90, doi:10.1111/j.1574-6968.2008.01409.x (2009).

27 Parkhill, J. et al. Genome sequence of Yersinia pestis, the causative agent of plague. Nature 413, 523–527, doi:10.1038/35097083 (2001).

28 Erickson, D. L., Jarrett, C. O., Wren, B. W. & Hinnebusch, B. J. Serotype differences and lack of biofilm formation characterize Yersinia pseudotuberculosis infection of the Xenopsylla cheopis flea vector of Yersinia pestis. J Bacteriol 188, 1113–1119, doi:10.1128/JB.188.3.1113-1119.2006 (2006).

29 Sun, F. et al. Fur is a repressor of biofilm formation in Yersinia pestis. PLoS One 7, e52392, doi:10.1371/journal.pone.0052392 (2012).

30 Nuorti, J. P. et al. A widespread outbreak of Yersinia pseudotuberculosis O:3 infection from iceberg lettuce. J Infect Dis 189, 766–774, doi:10.1086/381766 (2004).

31 Kangas, S. et al. Yersinia pseudotuberculosis O:1 traced to raw carrots, Finland. Emerg Infect Dis 14, 1959–1961, doi:10.3201/eid1412.080284 (2008).

32 Parn, T. et al. Outbreak of Yersinia pseudotuberculosis O:1 infection associated with raw milk consumption, Finland, spring 2014. Euro Surveill 20, doi:10.2807/1560-7917.ES.2015.20.40.30033 (2015).

33 Castro, H. et al. Genomic Epidemiology and Phenotyping Reveal on-Farm Persistence and Cold Adaptation of Raw Milk Outbreak-Associated Yersinia pseudotuberculosis. Front Microbiol 10, 1049, doi:10.3389/fmicb.2019.01049 (2019).

34 Jia, K. et al. Preliminary Transcriptome Analysis of Mature Biofilm and Planktonic Cells of Salmonella Enteritidis Exposure to Acid Stress. Front Microbiol 8, 1861, doi:10.3389/fmicb.2017.01861 (2017).

35 Tavares, E. R. et al. Phenotypic characteristics and transcriptome profile of Cryptococcus gattii biofilm. Sci Rep 9, 6438, doi:10.1038/s41598-019-42896-2 (2019).

36 Tan, X. et al. Transcriptome analysis of the biofilm formed by methicillin-susceptible Staphylococcus aureus. Sci Rep 5, 11997, doi:10.1038/srep11997 (2015).

37 Qin, N. et al. RNA-Seq-based transcriptome analysis of methicillin-resistant Staphylococcus aureus biofilm inhibition by ursolic acid and resveratrol. Sci Rep 4, 5467, doi:10.1038/srep05467 (2014).

38 Tram, G. et al. RNA Sequencing Data Sets Identifying Differentially Expressed Transcripts during Campylobacter jejuni Biofilm Formation. Microbiol Resour Announc 9, doi:10.1128/MRA.00982-19 (2020).

39 Jeske, A., Arce-Rodriguez, A., Thoming, J. G., Tomasch, J. & Haussler, S. Evolution of biofilm-adapted gene expression profiles in lasR-deficient clinical Pseudomonas aeruginosa isolates. NPJ Biofilms Microbiomes 8, 6, doi:10.1038/s41522-022-00268-1 (2022).

40 Tan, S. Y., Tan, I. K., Tan, M. F., Dutta, A. & Choo, S. W. Evolutionary study of Yersinia genomes deciphers emergence of human pathogenic species. Sci Rep 6, 36116, doi:10.1038/srep36116 (2016).

41 Choong, F. X. et al. Real-time optotracing of curli and cellulose in live Salmonella biofilms using luminescent oligothiophenes. NPJ Biofilms Microbiomes 2, 16024, doi:10.1038/npjbiofilms.2016.24 (2016).

42 Atkinson, S. et al. Biofilm development on Caenorhabditis elegans by Yersinia is facilitated by quorum sensing-dependent repression of type III secretion. PLoS Pathog 7, e1001250, doi:10.1371/journal.ppat.1001250 (2011).

43 Firoj Mahmud, A. K. M., Delhomme, N., Nandi, S. & Fallman, M. ProkSeq for complete analysis of RNA-Seq data from prokaryotes. Bioinformatics, doi:10.1093/bioinformatics/btaa1063 (2020).

44 Benda, M., Schulz, L. M., Stulke, J. & Rismondo, J. Influence of the ABC Transporter YtrBCDEF of Bacillus subtilis on Competence, Biofilm Formation and Cell Wall Thickness. Front Microbiol 12, 587035, doi:10.3389/fmicb.2021.587035 (2021).

45 Kostakioti, M., Hadjifrangiskou, M. & Hultgren, S. J. Bacterial biofilms: development, dispersal, and therapeutic strategies in the dawn of the postantibiotic era. Cold Spring Harb Perspect Med 3, a010306, doi:10.1101/cshperspect.a010306 (2013).

46 Fang, N. et al. RcsAB is a major repressor of Yersinia biofilm development through directly acting on hmsCDE, hmsT, and hmsHFRS. Sci Rep 5, 9566, doi:10.1038/srep09566 (2015).

47 Szklarczyk, D. et al. STRING v11: protein-protein association networks with increased coverage, supporting functional discovery in genome-wide experimental datasets. Nucleic Acids Res 47, D607–D613, doi:10.1093/nar/gky1131 (2019).

48 Merritt, M. E. & Donaldson, J. R. Effect of bile salts on the DNA and membrane integrity of enteric bacteria. J Med Microbiol 58, 1533–1541, doi:10.1099/jmm.0.014092-0 (2009).

49 Shah, M. K. & Bergholz, T. M. Variation in growth and evaluation of cross-protection in Listeria monocytogenes under salt and bile stress. J Appl Microbiol 129, 367–377, doi:10.1111/jam.14607 (2020).

50 Traxler, M. F. et al. The global, ppGpp-mediated stringent response to amino acid starvation in Escherichia coli. Mol Microbiol 68, 1128–1148, doi:10.1111/j.1365-2958.2008.06229.x (2008).

51 Abranches, J., Candella, M. M., Wen, Z. T., Baker, H. V. & Burne, R. A. Different roles of EIIABMan and EIIGlc in regulation of energy metabolism, biofilm development, and competence in Streptococcus mutans. J Bacteriol 188, 3748–3756, doi:10.1128/JB.00169-06 (2006).

52 Lazazzera, B. A. The phosphoenolpyruvate phosphotransferase system: as important for biofilm formation by Vibrio cholerae as it is for metabolism in Escherichia coli. J Bacteriol 192, 4083–4085, doi:10.1128/JB.00641-10 (2010).

53 Munoz-Elias, E. J., Marcano, J. & Camilli, A. Isolation of Streptococcus pneumoniae biofilm mutants and their characterization during nasopharyngeal colonization. Infect Immun 76, 5049–5061, doi:10.1128/IAI.00425-08 (2008).

54 Loo, C. Y., Mitrakul, K., Voss, I. B., Hughes, C. V. & Ganeshkumar, N. Involvement of an inducible fructose phosphotransferase operon in Streptococcus gordonii biofilm formation. J Bacteriol 185, 6241–6254, doi:10.1128/jb.185.21.6241-6254.2003 (2003).

55 Shime-Hattori, A. et al. Two type IV pili of Vibrio parahaemolyticus play different roles in biofilm formation. FEMS Microbiol Lett 264, 89–97, doi:10.1111/j.1574-6968.2006.00438.x (2006).

56 Jurcisek, J. A. et al. The PilA protein of non-typeable Haemophilus influenzae plays a role in biofilm formation, adherence to epithelial cells and colonization of the mammalian upper respiratory tract. Mol Microbiol 65, 1288–1299, doi:10.1111/j.1365-2958.2007.05864.x (2007).

57 Bahar, O., Goffer, T. & Burdman, S. Type IV Pili are required for virulence, twitching motility, and biofilm formation of acidovorax avenae subsp. Citrulli. Mol Plant Microbe Interact 22, 909–920, doi:10.1094/MPMI-22-8-0909 (2009).

58 Conrad, J. C. Physics of bacterial near-surface motility using flagella and type IV pili: implications for biofilm formation. Res Microbiol 163, 619–629, doi:10.1016/j.resmic.2012.10.016 (2012).

59 Burrows, L. L. Pseudomonas aeruginosa twitching motility: type IV pili in action. Annu Rev Microbiol 66, 493–520, doi:10.1146/annurev-micro-092611-150055 (2012).

60 Craig, L., Forest, K. T. & Maier, B. Type IV pili: dynamics, biophysics and functional consequences. Nat Rev Microbiol 17, 429–440, doi:10.1038/s41579-019-0195-4 (2019).

61 Pisithkul, T. et al. Metabolic Remodeling during Biofilm Development of Bacillus subtilis. mBio 10, doi:10.1128/mBio.00623-19 (2019).

62 Milton, D. L., Norqvist, A. & Wolf-Watz, H. Cloning of a metalloprotease gene involved in the virulence mechanism of Vibrio anguillarum. J Bacteriol 174, 7235–7244, doi:10.1128/jb.174.22.7235-7244.1992 (1992).

63 Merritt, J. H., Kadouri, D. E. & O’Toole, G. A. Growing and analyzing static biofilms. Curr Protoc Microbiol Chapter 1, Unit 1B 1, doi:10.1002/9780471729259.mc01b01s00 (2005).

64 Oehmcke, S. et al. A novel role for pro-coagulant microvesicles in the early host defense against streptococcus pyogenes. PLoS Pathog 9, e1003529, doi:10.1371/journal.ppat.1003529 (2013).

65 Patro, R., Duggal, G., Love, M. I., Irizarry, R. A. & Kingsford, C. Salmon provides fast and bias-aware quantification of transcript expression. Nature Methods 14, 417–419, doi:10.1038/nmeth.4197 (2017).

66 Risso, D., Ngai, J., Speed, T. P. & Dudoit, S. Normalization of RNA-seq data using factor analysis of control genes or samples. Nat Biotechnol 32, 896–902, doi:10.1038/nbt.2931 (2014).

