## Supplemental Fig S1,S2,S3 for "A core transcriptional response for biofilm formation by *Y. pseudotuberculosis*"

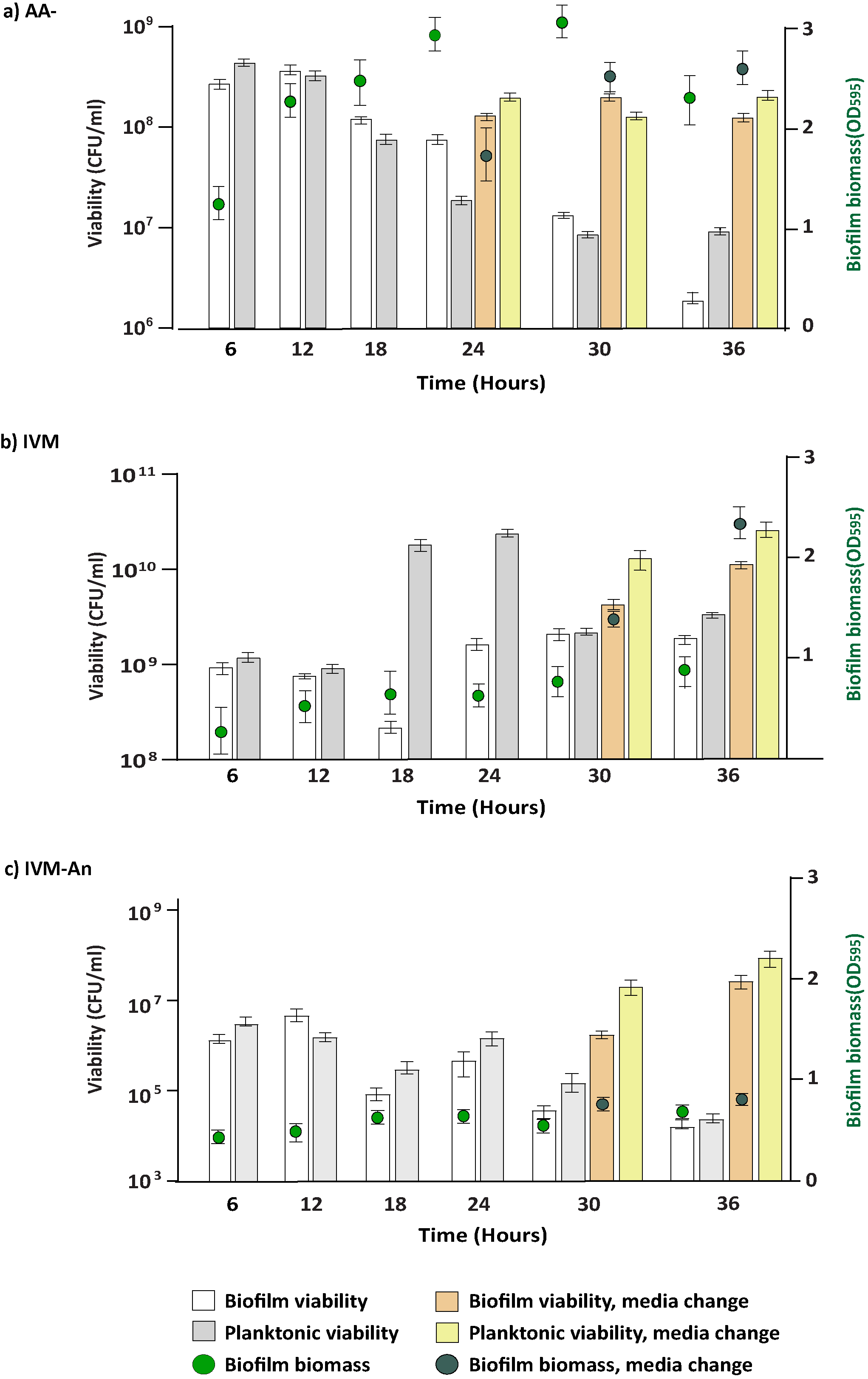

**Figure S1: Viability and Biofilm mass measurements with or without media change.** Viable counts of *Y. pseudotuberculosis* biofilm and planktonic bacteria under different conditions with or without change meadia at certain time point. White and grey bars) and corresponding biofilm biomass (green right axis and points) showing measurements without changing media. Light brown and yellow bars and corresponding biofilm biomass (dark green right axis and points) indicates measurement after media change. a) biofilm induced by AA-; media changed after 18 hours c) and d) biofilm formed in IVM and IVM-An; media changed after 24 hours. Three individual biological replicates were used for the viability and biofilm biomass measurement and presented as CFU per ml +/- SEM and OD_595_ +/- SEM.

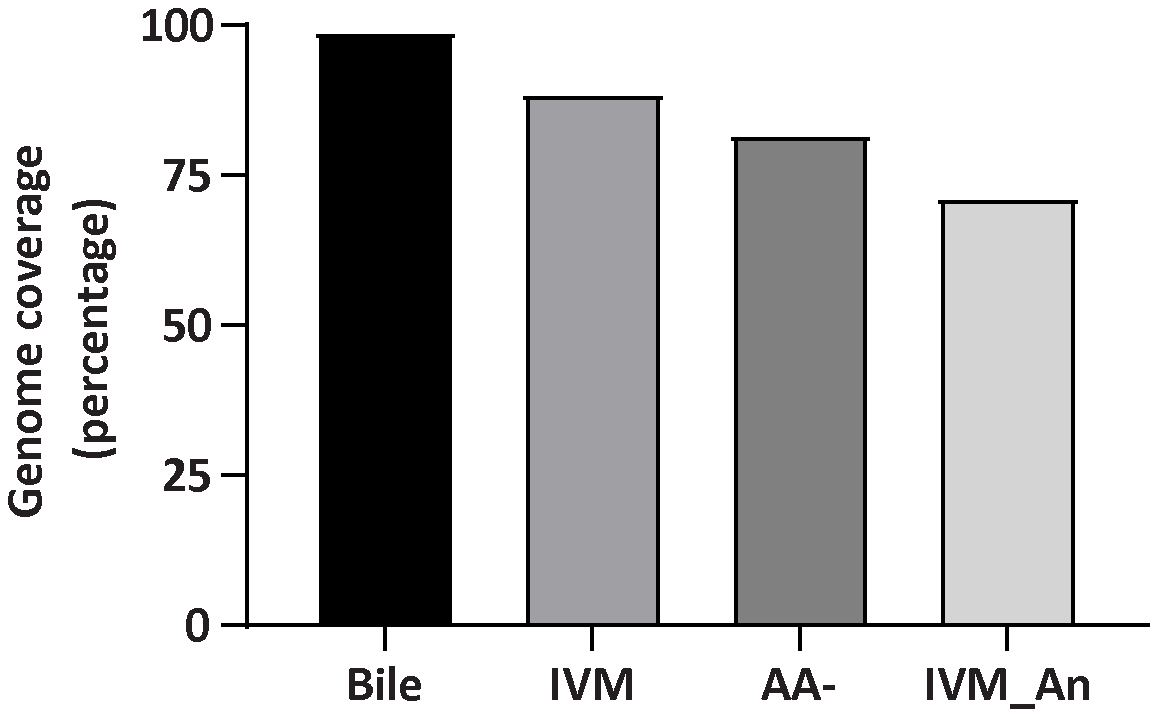

**Figure S2: Average cenome coverage of different biofilm RNA-Seq data.** Genome coverage is calculated after quality filtering and read processing of RNA-Seq data. Y axis denotes percentage of genes are covered by RNA-Seq data.

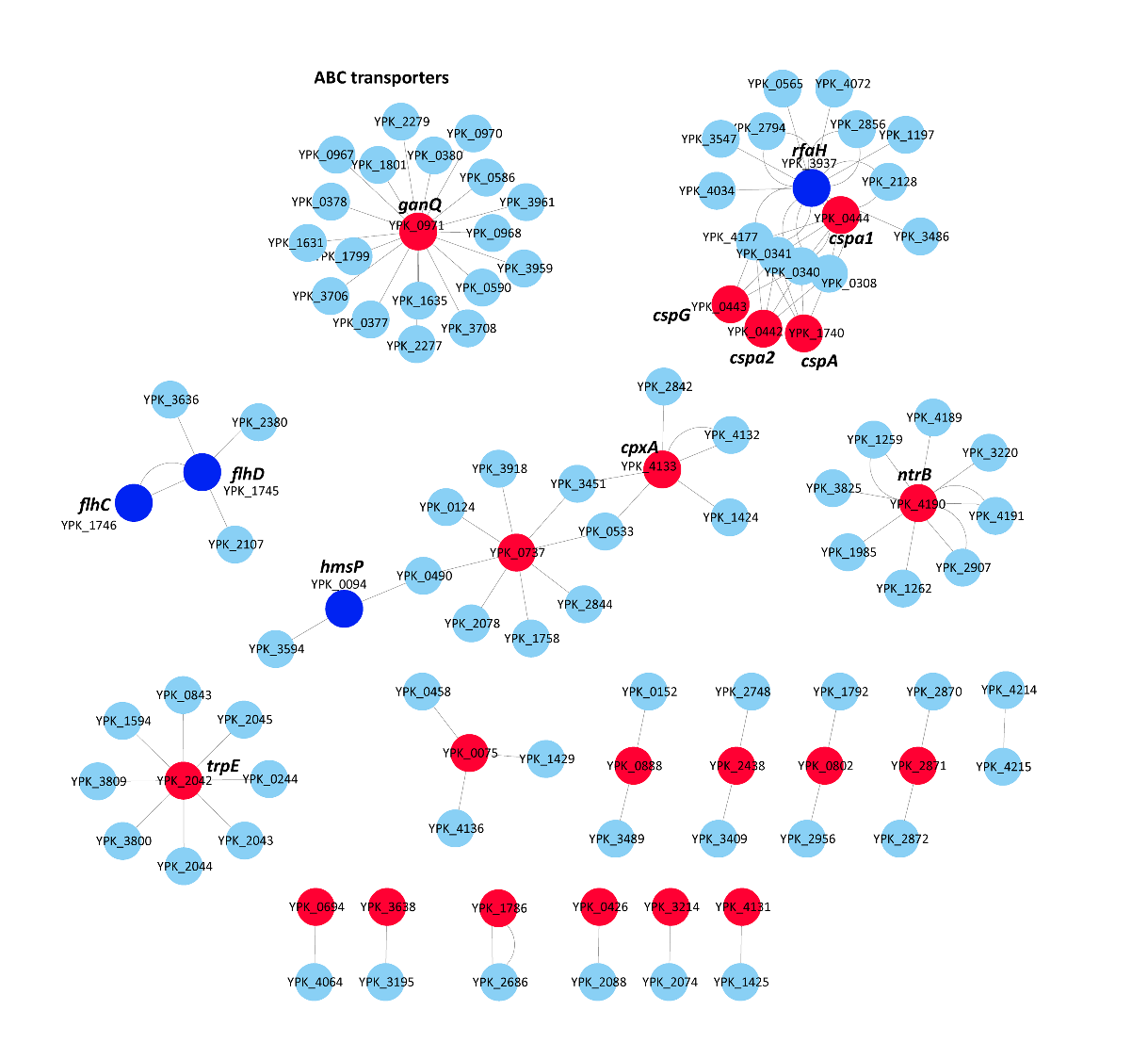

**Figure S3: Protein-protein interection network of mature biofilm core genes**. Experimentally validated protein interection data were collected from string database and Cytoscape (V. 3.6) sofatware was used to create the network. Red and blue color indicates the mature biofilm core genes while blue indicates downregulation and red indicates upregulation in biofilm
