## Supplemental Table S1,S4 for "A core transcriptional response for biofilm formation by *Y. pseudotuberculosis*"

| **LB *#** | **26**°**C** | **37**°**C** |
| --- | --- | --- |
| Control | + | ++ |
| 0.2% Glucose | - | - |
| 2.5 mM MgCl_2_ | + | + |
| 2.5 mM Ca^2+^ | + | + |
| 0.5% Bile | ++ | +++ |
| 25 µM Deferiprone | - | - |
| 0.1% Triton-X 100 | + | ++++ |
| **IVM *#** | **26°C** | **37°C** |
| Aerobic atmosphere (IVM) | - | ++++ |
| Anaerobic atmosphere (IVM-An) | + | +++ |
| **M9 *#** | **26**°**C** | **37**°**C** |
| Control | - | +++ |
| 1% Casamino acids | + | ++++ |
| **MOPS *#** | **26**°**C** | **37**°**C** |
| Control | + | +++ |
| Depletion of trp, thr, his | ++ | ++++ |

*summary from three biological and two technical experiments

\# incubation time 48 hours

**Table S2 and Table S3** : The supplementary table S2 and S3 can be downloaded from the following link http://www.fallmanlab.org/wp-content/uploads/2021/03/Supplementary-Table-S2-S3.zip

**Table S4**

| **Construct** | **Vector** | **Primer** | **Sequence (5' to 3')** |
| --- | --- | --- | --- |
| ∆YPK_0174 | pDM4 | 0174.A | accgtcgaccctcgaggttgtgccgatgccttagt |
|  |  | 0174.B | ggcttagatctttggcatgtcataactcctggatcgtc |
|  |  | 0174.C | aggagttatgacatgccaaagatctaagccagtgttag |
|  |  | 0174.D | tggaattcccgggagagctcgagagaacaggcagatcgta |
| ∆YPK_0444-0442 | pDM4 | 0444.A | accgtcgaccctcgagcatgcctacggactggtt |
|  |  | 0444.B | tgacggttacagcgctccaaaccgatgaagctatttaac |
|  |  | 0442.C | cttcatcggtttggagcgctgtaaccgtcaataat |
|  |  | 0442.D | tggaattcccgggagagctcccgcttttgatgatctccag |
| ∆YPK_0685 | pDM4 | 0685.A | accgtcgaccctcgagaacgtgatacctgacatgcc |
|  |  | 0685.B | cgcctgctttctacgggcgttagctgaattttcaaac |
|  |  | 0685.C | aattcagctaacgcccgtagaaagcaggcgaaaga |
|  |  | 0685.D | tggaattcccgggagagctcagccaatcgtacatattcgg |
| ∆YPK_0791 | pDM4 | 0791.A | accgtcgaccctcgagacgatgagagtctggttca |
|  |  | 0791.B | cagaatcggcttaatcattccctttctccatttactg |
|  |  | 0791.C | tggagaaagggaatgattaagccgattctggcataacg |
|  |  | 0791.D | tggaattcccgggagagctcgaatcgagccgtttcttcgt |
| ∆YPK_0971 | pDM4 | 0971.A | accgtcgaccctcgaggagtaccagcaccaaattgc |
|  |  | 0971.B | agtacttcctgacgtcatgttcccgtcccttatg |
|  |  | 0971.C | agggacgggaacatgacgtcaggaagtactaaggg |
|  |  | 0971.D | tggaattcccgggagagctcgctttcgcacgtttagcca |
| ∆YPK_1186 | pDM4 | 1186.A | accgtcgaccctcgagcctctacaaaggctgtattcc |
|  |  | 1186.B | ggcatgataagcaggcataatactctcctcgttaccg |
|  |  | 1186.C | gaggagagtattatgcctgcttatcatgcctaatca |
|  |  | 1186.D | tggaattcccgggagagctctgagaccgaccatcaattac |
| ∆YPK_1428 | pDM4 | 1428.A | accgtcgaccctcgaggcagtacaacaaaagatggcatg |
|  |  | 1428.B | cgttgattactcaagctggaacattgtgtttccccaa |
|  |  | 1428.C | aacacaatgttccagcttgagtaatcaacgcgctctt |
|  |  | 1428.D | tggaattcccgggagagctctcagcagaggcatgctcat |
| ∆YPK_1449 | pDM4 | 1449.A | accgtcgaccctcgaggtttaaatcgcaacccataag |
|  |  | 1449.B | ggtttattgcgcgggagcggttataaacatcgtgct |
|  |  | 1449.C | atgtttataaccgctcccgcgcaataaaccttaatg |
|  |  | 1449.D | tggaattcccgggagagctcatccataaacggagcagact |
| ∆YPK_1705 | pDM4 | 1705.A | accgtcgaccctcgagccaagacgaccatagttgaag |
|  |  | 1705.B | tggtattacgctcggggctaacattgatttgttcacagg |
|  |  | 1705.C | aaatcaatgttagccccgagcgtaataccaaatctgtt |
|  |  | 1705.D | tggaattcccgggagagctcgatttaccgtttggaagggc |
| ∆YPK_1745 | pDM4 | 1745.A | accgtcgaccctcgaggagtagaattggtcctctttagg |
|  |  | 1745.B | accatcaatcatgctcgtactcatcttacacatccca |
|  |  | 1745.C | tgtaagatgagtacgagcatgattgatggttgagaaaag |
|  |  | 1745.D | tggaattcccgggagagctcaacatgccactgtcaacgaa |
| ∆YPK_2042 | pDM4 | 2042.A | accgtcgaccctcgaggcttacaagacgcgaacgtt |
|  |  | 2042.B | ccattagaaaatctcacgtgatgtttgcatcattgag |
|  |  | 2042.C | atgcaaacatcacgtgagattttctaatggccgatatcc |
|  |  | 2042.D | tggaattcccgggagagctcacataccttgcccatcatgt |
| ∆YPK_2223 | pDM4 | 2223.A | accgtcgaccctcgagtactaccaggtgcctactac |
|  |  | 2223.B | ggtgtgctatcaacgattcatgggcgtacttctcc |
|  |  | 2223.C | agtacgcccatgaatcgttgatagcacacctattacttg |
|  |  | 2223.D | tggaattcccgggagagctcatagcgctcaggtattgcag |
| ∆YPK_3337 | pDM4 | 3337.A | accgtcgaccctcgagcatcgcctcaatctgtgcat |
|  |  | 3337.B | gaattactgttccgaggagaatgaactttccattatgagg |
|  |  | 3337.C | gaaagttcattctcctcggaacagtaattccactatttc |
|  |  | 3337.D | tggaattcccgggagagctcccgtcattttgcacgttgta |
| ∆YPK_3368, luxS | pDM4 | luxS.A | accgtcgaccctcgagtgagttggctgagcattatc |
|  |  | luxS.B | tgatactgagcactatggcatttagttacctcctca |
|  |  | luxS.C | ggtaactaaatgccatagtgctcagtatcagtaggc |
|  |  | luxS.D | tggaattcccgggagagctcgaagttaatccacagagcga |
| ∆YPK_3425, rpoS | pDM4 | rpoS.A | accgtcgaccctcgaggagagattggcggtatcttatga |
|  |  | rpoS.B | cagcatatgagccaattccgcgaataagtgtgtc |
|  |  | rpoS.C | cacttattcgcggaattggctcatatgctgctc |
|  |  | rpoS.D | tggaattcccgggagagctccgatagcttctctgcttcc |
| ∆YPK_3616 | pDM4 | 3616.A | accgtcgaccctcgagcacatccggttcaaactcaacg |
|  |  | 3616.B | gcatgtcacctatcccgtgttcatcgtgcattagc |
|  |  | 3616.C | tgcacgatgaacacgggataggtgacatgcagaaaatgc |
|  |  | 3616.D | tggaattcccgggagagctcagtcggtatgaccatcaacc |
| ∆YPK_3648 | pDM4 | 3648.A | accgtcgaccctcgaggcggatgtgattgtcgca |
|  |  | 3648.B | ttagaggccaggagccattgcgcgctcctcttg |
|  |  | 3648.C | gaggagcgcgcaatggctcctggcctctaaaatgg |
|  |  | 3648.D | tggaattcccgggagagctcggcaggtgatgctgtcttat |
| ∆YPK_3937, rfaH | pDM4 | 3937.A | gccgctcgagctgttcca gtaatgagag |
|  |  | 3937.B | gtaagtcagcgtgttaagtttcacatacctttgg |
|  |  | 3937.C | taacacgctgacttacaataatc |
|  |  | 3937.D | ctcttctagacctatcaggtgtctgg |
| ∆YPK_4131 | pDM4 | 4131.A | accgtcgaccctcgaggtctgatggtgttggcgta |
|  |  | 4131.B | tacttacttctgggcacgcatcgttaactcctaaagttc |
|  |  | 4131.C | gagttaacgatgcgtgcccagaagtaagtagtagtac |
|  |  | 4131.D | tggaattcccgggagagctccatcattgcactccagcaaa |
| ∆YPK_4190 | pDM4 | 4190.A | accgtcgaccctcgaggctatcgatgcgtatatcgagc |
|  |  | 4190.B | tgcatagaaacctcacataatgcagactcctgcacag |
|  |  | 4190.C | ggagtctgcattatgtgaggtttctatgcaacgagg |
|  |  | 4190.D | tggaattcccgggagagctcatagcccgttcaaccagag |
